# Temperature-induced reorganisation of Schistocephalus solidus (Cestoda) proteome during the transition to the warm-blooded host

**DOI:** 10.1101/2021.04.08.439102

**Authors:** Ekaterina V. Borvinskaya, Albina A. Kochneva, Polina B. Drozdova, Olga V. Balan, Victor G. Zgoda

## Abstract

The protein composition (proteome) of cestode *Schistocephalus solidus* was measured in an experiment simulating the transition of the parasite from a cold-blooded to a warm-blooded host. Infective *S. solidus* plerocercoids obtained from the three-spined stickleback *Gasterosteus aculeatus* were heated at 40 °C for 1 hour or cultured *in vitro* at 40 °C and 22 °C for 48 hours. In the short-term experiment, the content of only one tegument protein decreased after heating. After long-term heating, which triggered parasite sexual maturation, an increase in the content of ribosomal proteins, translation initiation factors and enzymes of the amino acid biosynthesis pathway was observed. The synthesis of certain gene products for carbohydrate metabolism, including glycolysis/gluconeogenesis, was found to be regulated in the parasite by temperature.

**Summary statement:** The study focuses on the processes that determine the survival of parasites in warm-blooded animals, and thus demonstrates the potential for the application of proteomics in veterinary medicine.

## Introduction

Cestodes are obligate parasites with a complex life cycle, during which they are trophically transmitted from one type of host to another, often undergoing dramatic changes in environmental conditions. Moreover, the stress associated with the new environment in the new host even stimulates metamorphosis or growth of the parasites. To explain the amazing ecological plasticity of these parasites, it is of interest to study the mechanisms of rapid and slow adaptation of cestodes, especially the molecular composition and gene regulation of tapeworms in response to external stimuli.

The parasitic worm *Schistocephalus solidus* (Müller, 1776), due to its specific ontogeny, is a suitable organism for studying the transition of cestodes from cold-blooded to warm-blooded hosts. The first intermediate host of *S. solidus* is a freshwater copepod, while the second is the cold-water three-spined stickleback *Gasterosteus aculeatus* (Linnaeus, 1758), which receives the parasite when it feeds on infected zooplankton. In the body cavity of fish, helminths develop into the second larval stage (plerocercoid), which actively grows and accumulates nutrients. A plerocercoid that has reached a weight of about 0.5 g becomes infective, that is able to survive in the final hosts, piscivorous birds (Tierney and Crompton, 1992). If swallowed by a bird with a host fish, the parasite in the avian intestines undergoes 20-40 °C heating in several minutes. This sharp rise in the ambient temperature triggers an approximately 36-hour maturation and fertilization program of the parasite, which ends with the adult worm releasing its eggs and leaving the host (Smyth, 1950; Hopkins and Smyth, 1951).

The transcription of *S. solidus* genes, which presumably provide the successful colonization of a warm-blooded host, have been previously investigated by Hébert with colleagues in the model *in vitro* experiment (Hébert et al., 2017). Overexpression of certain gene products has been reported either in larvae or in adult worms (the “molecular signature” of the corresponding life stage). Gravid worms have been shown to possess enhanced pathways of reproduction and redox homoeostasis compared to infectious plerocercoids. Considering the possible differences between the actual *S. solidus* protein and transcript profiles due to RNA and protein processing, it would be useful to confirm the transcriptomic data by direct measurement of the parasite proteins. It would also be interesting to investigate the molecular rearrangements that occur in the very first moments after the temperature rise, as the key events that triggers the maturation of *S. solidus* in the final host.

Therefore, the aim of this study was to evaluate how the composition of proteins of *S. solidus* changes in an experiment simulating the transition of the parasite from fish to warm-blooded birds during short-term (1 hour) and long-term (48 hours) incubation of infectious plerocercoids at 40 °C.

## Materials & Methods

### Animal collection

Individuals of the three-spined stickleback *G. aculeatus* infected with *S. solidus* were caught with an aquarium net in August 2018 and 2019 in the “Lake Mashinoe” (North-West of Russia, 66°17’46.5”N 33°21’59.3”E). Infected fish were easily distinguished by their behaviour (swimming near the water surface) and a swollen abdomen. Live fish were delivered to the laboratory in barrels with water (with aeration) at 20-22 °C and kept under such conditions until the experiment (no more than five days).

### Incubation of S. solidus plerocercoids

#### Short-term heating (1 hour)

Three-spined sticklebacks were killed by destroying the brain with a needle. The fish were then individually tightly wrapped with cling film to prevent the plerocercoids from crawling out of the host’s body cavity and contacting the environment during incubation. Thereafter, the infected sticklebacks (n=3) were immediately placed in water heated to 40 °C for one hour (Fig. 1). At the end of incubation, the fish were quickly taken out from the water, *S. solidus* were removed from the host body cavity, weighed and frozen in liquid nitrogen. Plerocercoids taken out from the body cavity of sticklebacks immediately after their sacrifice (n=3) were used as a control group. This experiment was repeated twice, in 2018 and 2019.

**Figure 1.**
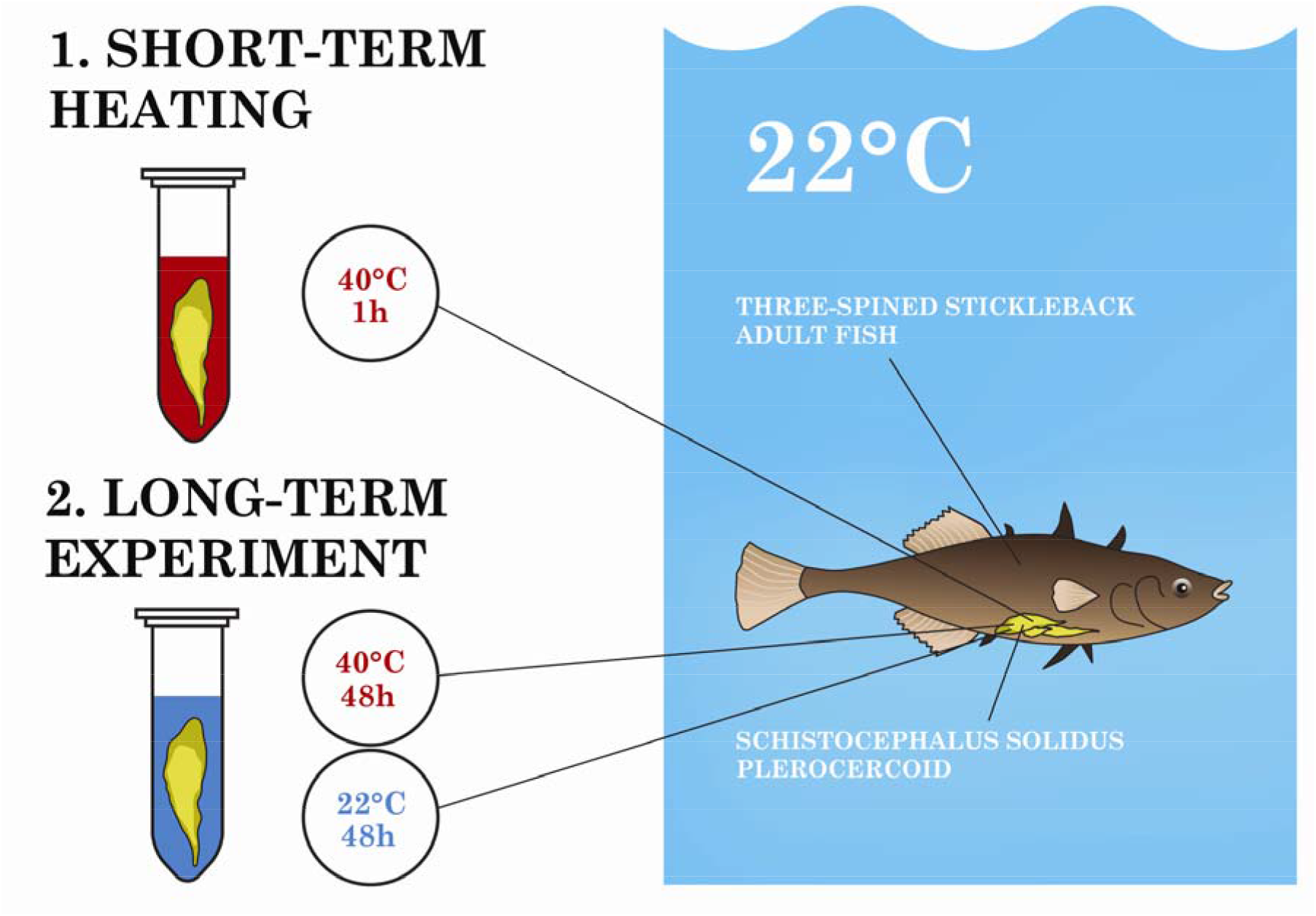
Experimental design.

### Long-term heating (48 hours, heat-induced maturation)

The infected sticklebacks were killed by the destruction of the brain using a needle, while preventing pressure on the abdomen of the fish. The worms were removed from the fish body cavity and washed two times in a solution containing RPMI-1640 medium (Sigma-Aldrich) with 1% antibiotic antimycotic solution (Sigma-Aldrich). Next, the worms were placed in culture flasks with culture medium (RPMI-1640 medium, 0.1% antibiotic antimycotic solution and 10% glucose) and incubated in the water jacketed incubator (SHEL LAB) at 40 °C (n=5) and 20-22 °C (n=5) in a 5-10% CO_2_ atmosphere.

Egg production was assessed visually twice a day. In the treatment group the parasites became mature within 48 hours, as determined by the presence of eggs in the medium. After this the helminths from both groups were removed from the culture flasks, washed in fresh culture medium, frozen in liquid nitrogen and stored until proteome analysis. Plerocercoids taken out from the body cavity of sticklebacks immediately after their sacrifice (n=3) were used as a reference group. The whole long-term experiment was carried out in 2019.

### Sample preparation and LC-MS/MS analysis of S. solidus proteins

Frozen helminths were ground with a pestle in a mortar with liquid nitrogen to a powder with the addition of 0.1 mol/L Tris-HCl (pH 7.6) with 1% protease inhibitor cocktail (Amresko) and 1% phenylmethanesulfonyl fluoride (Sigma-Aldrich). After thawing the extraction buffer protein was precipitated with 100% trichloroacetic acid (final concentration 10%)(Sigma-Aldrich). After centrifugation at 12000 g for 5 min the pellet was washed with ice-cold 80% ethanol and ice-cold acetone by successive centrifugation-resuspension cycles. The final samples were lyophilized (Labconco FreeZone 6L) and stored until analysis at −80 °C.

The lyophilisates were resuspended in the extraction buffer (300 μL) containing 4% SDS (Sigma-Aldrich) and 0.1 mol/L 1,4-dithiothreitol (Roche) in 0.1 mol/L Tris-HCl (pH 7.6). The total protein content in samples was measured according to the BCA method (*Basic Protein* …, 1994). A total protein amount of 100 mg for each sample was used for tryptic digestion according to the common FASP protocol (Wiśniewski et al., 2009). Briefly, protein disulfide bridges were reduced with 100 mmol/L 1,4-dithiothreitol in 100 mmol/L Tris-HCl (pH 8.5), and alkylation of thiols was performed with 55 mmol/L iodoacetamide (Sigma-Aldrich) in 8 mol/L urea in 100 mM Tris-HCl (pH 8.5). Detergents in the samples were exchanged with 100 mmol/L Tris-HCl (pH 8.5) using Microcon filters (10 kDa cut off, Millipore). Tryptic digestion with trypsin (Sequencing Grade Modified, Promega) to protein ratio of 1:100 was carried out overnight at 37 °C in a 50 mM tetraethylammonium bicarbonate (pH 8.5). To obtain the peptide solution the filter samples were centrifuged at 11000 g for 15 min at 20 °C. The filters were then washed with 50 mL of 30% formic acid solution (Sigma-Aldrich) by centrifugation at 11000 g for 15 min in a thermostatic centrifuge at 20 °C. The filtrates were dried in a vacuum concentrator and dissolved in 20 ml of 5% formic acid for subsequent LC-MS analysis.

The separation of peptide mixture was performed using an Ultimate 3000 RSLCnano chromatographic HPLC system (Thermo Scientific, USA) connected to a Q-exactive HFX mass spectrometer (Thermo Scientific, USA). One microgram of peptides in a volume of 1-4 µL was loaded onto the Acclaim µ-Precolumn (0.5 mm х 3 mm, 5 µm particle size, Thermo Scientific) at a flow rate of 10 µL/min for 4 min in an isocratic mode of Mobile Phase C (2% acetonitrile (Sigma-Aldrich), 0.1% formic acid). Then the peptides were separated with high-performance liquid chromatography (HPLC, Ultimate 3000 Nano LC System, Thermo Scientific, Rockwell, IL, USA) in a 15-cm long C18 column (Acclaim^®^ PepMap™ RSLC, inner diameter of 75 μm, Thermo Fisher Scientific, Rockwell, IL, USA). The peptides were eluted with a gradient of buffer B (80% acetonitrile, 0.1% formic acid) at a flow rate of 0.3 μL/min. The total run time was 90 minutes, which included initial 4 min of column equilibration to buffer A (0.1% formic acid), then gradient from 5 to 35% of buffer B over 65 min, then 6 min to reach 99% of buffer B, flushing 10 min with 99% of buffer B and 5 min re-equilibration to buffer A.

MS analysis of the samples was performed at least in triplicate with a Q Exactive HF-X mass spectrometer (Q Exactive HF-X Hybrid Quadrupole-OrbitrapTM Mass spectrometer, Thermo Fisher Scientific). The temperature of the capillary was 240 °C, and the voltage at the emitter was 2.1 kV. Mass spectra were acquired at a resolution of 120000 (MS) in a range of 300−1500 *m/z*. Tandem mass spectra of fragments were acquired at a resolution of 15000 (MS/MS) in the range from 100 *m/z* to m/z value determined by a charge state of the precursor, but no more than 2000 *m/z*. The maximum integration time was 50 ms and 110 ms for precursor and fragment ions, correspondingly. AGC target for precursor and fragment ions were set to 1^10^6^ and 2^10^5^, correspondingly. An isolation intensity threshold of 50000 counts was determined for precursor selection, and up to top 20 precursors were chosen for fragmentation with high-energy collisional dissociation (HCD) at 29 NCE. Precursors with a charge state of +1 and more than +5 were rejected and all measured precursors were dynamically excluded from triggering a subsequent MS/MS for 70 s.

### Protein quantification and functional annotation

The mass spectra raw-files were loaded into the MaxQuant v.1.6.4.3 program (Cox and Mann, 2008). The searches were performed using the Andromeda algorithm (built into MaxQuant) using the *Schistocephalus solidus* database loaded from UniProt/Swiss-Prot database (Proteome IDs UP000275846 and UP000050788 (Fontenla et al., 2017; International Helminth …, 2019, access in May 2020, 43048 sequences). The following search parameters were set: digestion enzyme was trypsin with a maximum of 2 missed cleavages; 5.0 ppm as MS1 and 0.01 Da as MS2 tolerances; fixed modification: carbamidomethylation (Cys); variable modifications: N-terminal proteins acetylation, and methionine oxidation (Met). Peptide Spectrum Matches (PSMs), peptides and proteins were validated at a 1% False Discovery Rate estimated using the decoy hit distribution.

Protein quantification was based on the label-free quantification (LFQ) method. The data matrix resulting from the MaxQuant analysis was loaded into Perseus software v. 1.6.2.3. The data was filtered to exclude proteins identified by modified (only identified by side) and reverse peptides, potential contaminants and proteins with fewer than two unique peptides. Obtained LFQ intensities performed in Table S1. An imputation of missing values procedure was performed for proteins that had LFQ-intensities for at least two from three technical repeats (Välikangas et al., 2018). Imputation of the missing values was done using the bpca method in the PCAmethod package for RStudio (Stacklies et al., 2007). The obtained data matrix of averaged replicates can be found in Table S2.

The functional annotation of identified *S. solidus* proteins was analysed using the Blast2GO, InterPro, BlastKOALA and QuickGO services (Götz et al., 2008; Binns et al., 2009; Kanehisa et al., 2016; Mitchell et al., 2019). Visualisation of KEGG pathway enrichment integrated with lipid enrichment was performed using the ‘‘Pathview’’ package (Luo and Brouwer, 2013) for the R computing environment. GO-term enrichment was analysed using the ‘‘topGO’’ package for R (Alexa and Rahnenfuhrer, 2010).

The presence of transmembrane regions was predicted using the TMHMM algorithm (version 2.0) and Phobius (Krogh et al., 2001; Brown et al., 2012). Signal peptides of classically and non-classicaly secreted proteins were predicted using SecretomeP 2.0 and SignalP 5.0 services (Bendtsen et al., 2004; Almagro Armenteros et al., 2019).

The unique *S. solidus* protein sequences were filtered from *S. solidus* protein database loaded from UniProt/Swiss-Prot (access in May 2020; 43048 sequence identifiers) using the CD-HIT Suite service with 100% sequence identity cut-off settings (Huang et al., 2010). The resulting groups, consisting of synonymous names of protein sequences, were used to compare the expression of *S. solidus* proteins with the previously described transcriptomic (Hébert et al., 2017) and other published data (Tables S3, S4).

### Statistical analysis

The comparison of protein expression in the control and experimental groups was performed with the ROTS package using a reproducibility-optimized test statistic adjusts a modified t-statistic (Suomi et al., 2017). Before analysis the LFQ-intensities were normalized using the normalizeVSN function in the limma package (Ritchie et al., 2015).

### Ethical approval

All animal experiments were in accordance with the EU Directive 2010/63/EU for animal experiments and the Declaration of Helsinki and was approved by the Animal Subjects Research Committee of the Institute of Biology at Irkutsk State University (Protocol *№*5, 16 Apryl 2018).

## Results

### Changes in protein profiles of S. solidus infective plerocercoids after short-term heating

The proteomic data obtained for worms exposed to 40 °C for 1 hour under semi-anaerobic conditions (n=3) were compared with those of infective plerocercoids taken directly from fish (n=3). This experiment was repeated twice, in 2018 and 2019. The numbers of proteins identified in the extracts of heated and non-heated plerocercoids were 514 and 402 in 2018 and 2019, respectively; these two sets of proteins overlap by 281 proteins (Table S3).

Multidimensional scaling analysis demonstrated a significant variability of the obtained data depending on the year of the experiment. This shift was probably caused by variations in the conditions of the chromatographic separation of the protein extract, despite following the same research protocol (Fig. S1). Thus, these two datasets were not combined, but analysed for differential expression separately. In the 2018 and 2019, the content of 151 and 13 proteins were differentially expressed in heated worms compared with not heated, respectively. Only one protein, A0A0X3PQ89, was identified as differentially expressed (down-regulated upon heating) in both data sets. Thus, only the degradation of the A0A0X3PQ89 tegument protein was confirmed by repeated measurements in *S. solidus* plerocercoids during the first hour of heating.

### Changes in protein profiles of S. solidus infective plerocercoids after long-term heating

In a long-term experiment, after 48 hours of incubation of *S. solidus* plerocercoids at 40 °C, the worms began to release eggs into the media, indicating their maturation and completion of fertilization. In the reference group, in which the worms were incubated at 22 °C in the same environment, no genital development and egg production was observed.

Since the stimulation of maturation was carried out in 2019, the proteomic composition of mature worms was compared with the proteome of infective plerocercoids obtained directly from fish in an experiment of the same year. After analysing the mass-spectra of the incubated worms, 1361 *S. solidus* proteins were identified. Of these, 662 proteins of heated worms and 434 proteins of worms incubated at 22 °C differed significantly in content compared to infectious plerocercoids from the fish body cavity (Table S4). The worms incubated at different temperatures shared 405 proteins with differential expression (Fig. 2A), and the concentration of all of them changed in the same direction.

**Figure 2.**
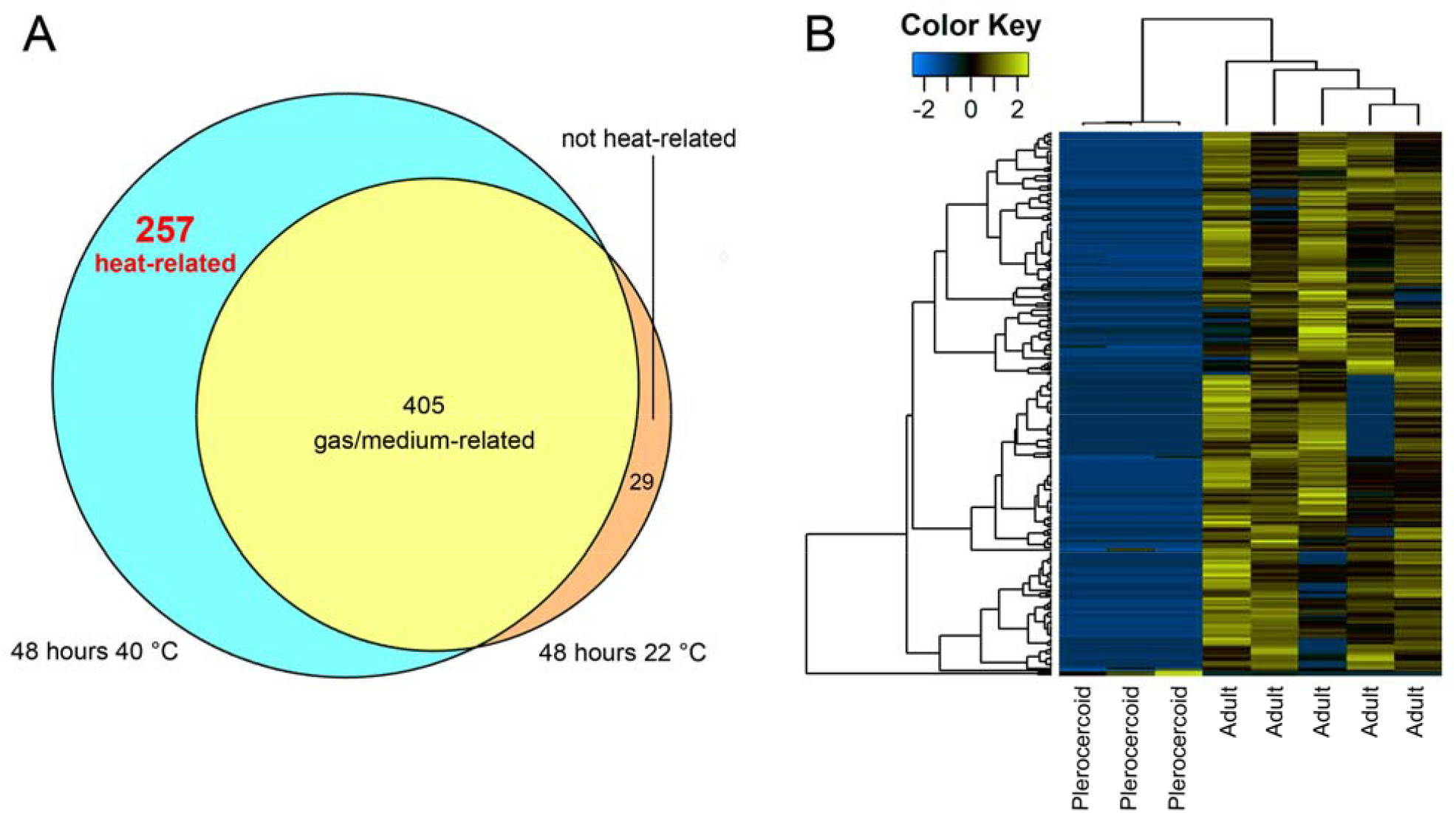
Differential expression of *S. solidus* infective plerocercoids and adult worms. (A) Venn diagrams represent the overlap of the lists of IDs of proteins differentially expressed between mature worms and infective plerocercoids. Blue indicates the number of proteins differentially expressed only in parasites incubated at 40 °C for 48 hours (“heat-related”), while yellow indicates proteins differentially expressed only in parasites incubated at 22 °C for 48 hours (“not heat-related”), and green indicates proteins differentially expressed in both experiments (“gas/medium-related). (B) Heatmap of *S. solidus* “heat-related” proteins, the level of which significantly changed only in worms that reached sexual maturity after incubation at 40 °C for 48 hours (for 5 biological replicates) in comparison with infective plerocercoids (for 3 biological replicates). The experiment was repeated twice.

The synthesis of 99% of proteins, the content of which changes only after incubation at 40 °C (these proteins are hereinafter referred as “heat-related”) and 98% of proteins, the content of which changes after incubation at both 40 °C and 22 °C (“gas/medium-related”), increased in comparison with these proteins in plerocercoids taken out from fish host (Fig. 2B, Table S4).

## Discussion

During the rapid adaptation to a sharp increase in temperature in the final host, a decrease in the content of the platyhelminth-specific Tegument-Allergen-Like protein A0A0X3PQ89 (TAL, antigen Sm21.7) was observed in the *S. solidus* plerocercoid after 1 hour of exposure at 40 °C under semi-anaerobic conditions (Table S3). Then, after 48 hours of heating, the level of A0A0X3PQ89 in gravid *S. solidus* again becomes comparable to that in infective larvae (Table S4). TALs are known to be involved in the organization and functioning of the tegument, but their biological role depends on the worm species and protein isoform (Fitzsimmons et al., 2012; Braschi et al., 2006; Zheng et al., 2013). For instance, all 13 members of *Schistosoma mansoni* TALs were shown to have different localization and patterns of expression in adult and larvae parasites (Fitzsimmons et al., 2012). In *Echinococcus multilocularis*, similar to *S. solidus*, A0A0X3PQ89 orthologue down-regulates then metacestode develop into pre-gravid adult worm, but gravid parasite again has high expression of the tegument protein (Zheng et al., 2013). We speculate that the initial decline in the content of tegument protein followed by recovery during worm maturation could be associated with tegument renovation occurring in the definitive host. At least in *S. solidus* plerocercoids the initiation of massive cuticular peeling was described within an hour after ambient temperature increased to 40 °C (Smyth, 1946). To elucidate the role of A0A0X3PQ89 in tegument loss or other adaptations at the early stages of final host colonization, further study of the localization of this protein is required.

The temperature increase triggers a maturation program encoded in the genomes of parasites, which launches a global restructuring of the protein profile and, as a result, changes the phenotype of the worm. In our long-term experiment, the infective *S. solidus* plerocercoids reached sexual maturity on the second day of exposure to the temperature of warm-blooded hosts, while the worms incubated at 22 °C under the same conditions remained immature during this time. This is consistent with physiological and histological studies showing that temperature rise is a necessary factor triggering *S. solidus* meiosis and reproductive behaviour (Hopkins, 1950; Smyth, 1952; Schjørring, 2003).

It was found that about six hundred proteins in worms incubated *in vitro* for 2 days have a concentration different from that of infectious plerocercoids from the body cavity of fish. Of these, up to two-thirds change both at 40 °C and 22 °C (Fig. 2A), indicating that their synthesis is under the control of the composition of the culture medium (’gas/medium-related”). These proteins changed synchronously, i.e., they increased or decreased in content in both experiments (Table S4). Since the incubation medium does not completely imitate liquid medium neither of the abdominal cavity of fish nor the intestines of birds, proteins from this group were not considered by us as proteins of interest for study of natural *S. solidus* proteome. For example, bile acids and lipid components of the chyme are known to affect the metabolism of cestodes (Frayha and Smyth, 1983), but the culture medium used in this experiment was depleted in these compounds. Another 257 proteins were differentially expressed only in worms incubated at higher temperature (’heat-related”), and they were in the focus of this study, as definitely associated with the adaptation of the parasite to a final host.

In our study, the number of proteins that were different between adult *S. solidus* and infectious plerocercoids was fewer than the number of differentially expressed genes in a similar experiment provided by Hébert and colleagues (2017). This can be explained by the lower sensitivity of proteomic methods compared to the analysis of the transcriptome, as well as the different effect of mRNA and/or protein processing in cells on the results of these two methods (Sun et al., 2014). It should also be noted that the study by Hébert and colleagues there were no reference group to evaluate the effect of culture media on the expression of proteins, which may have another transcription pattern in the avian intestine. Nevertheless, a comparison of independent transcriptomic and proteomic datasets revealed gene products that were altered in the same way in adult worms in both studies. From these, we filtered out 80 “heat-related” genes whose expression is regulated by temperature and which can be considered as reliable markers of the adult or plerocercoid stage (Table 1).

**Table 1.**
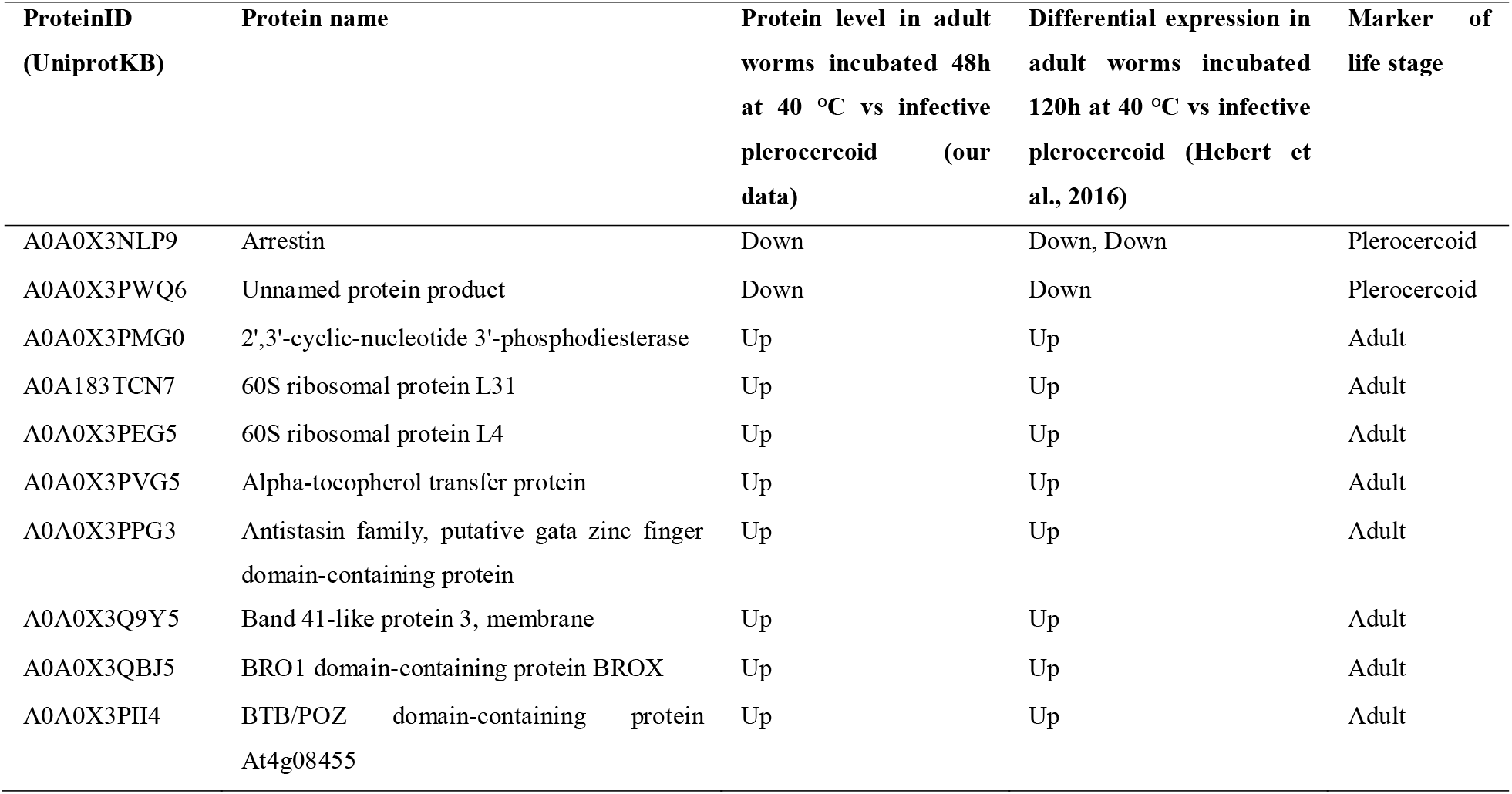
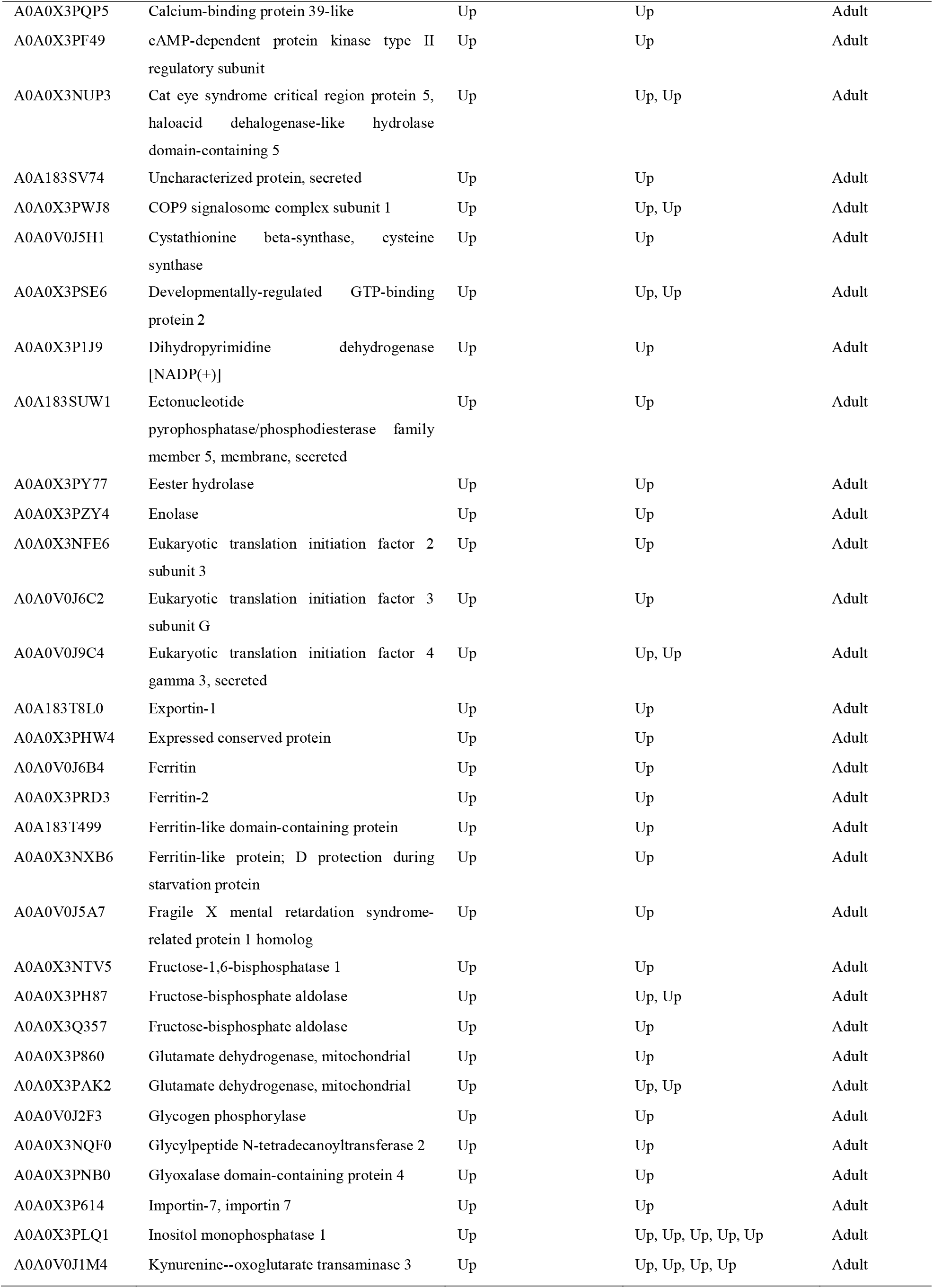
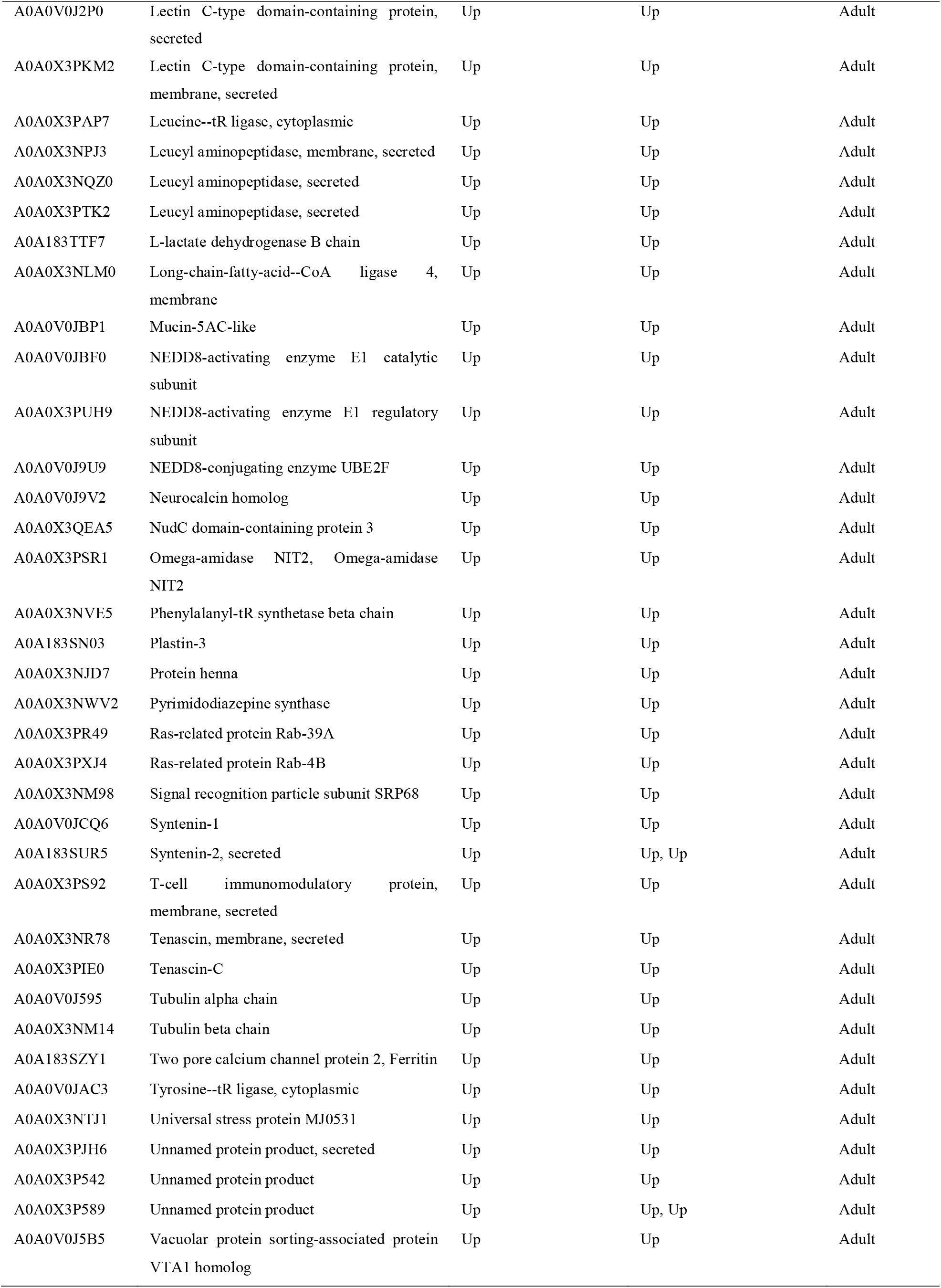
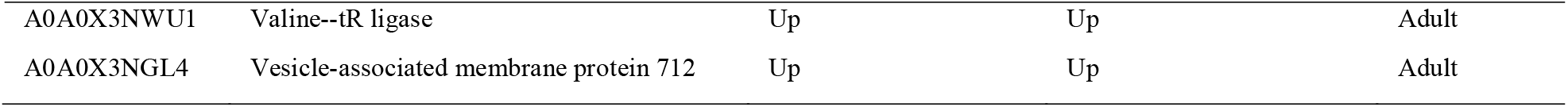
Differentially expressed “heat-related” proteins and corresponding transcripts in *S. solidus* infective plerocercoids and adult worms according to proteome (our data) and transcriptome studies (Hebert et al., 2016).

According to the obtained proteomic data, the content of almost all heat-related proteins is increased in adult worms as compared to plerocercoids. This indicates the global activation of the metabolism in maturing *S. solidus* after the dormant stage due to the need to complete the ontogenetic cycle in just a few days and convert the energy accumulated by the larvae into the maximum number of units of gametes and eggs. An intense rearrangement of the metabolism of adult parasites can also be seen due to the significant proportion of proteins involved in the processing of genetic information among heat-induced proteins (Fig. 3A).

**Figure 3.**
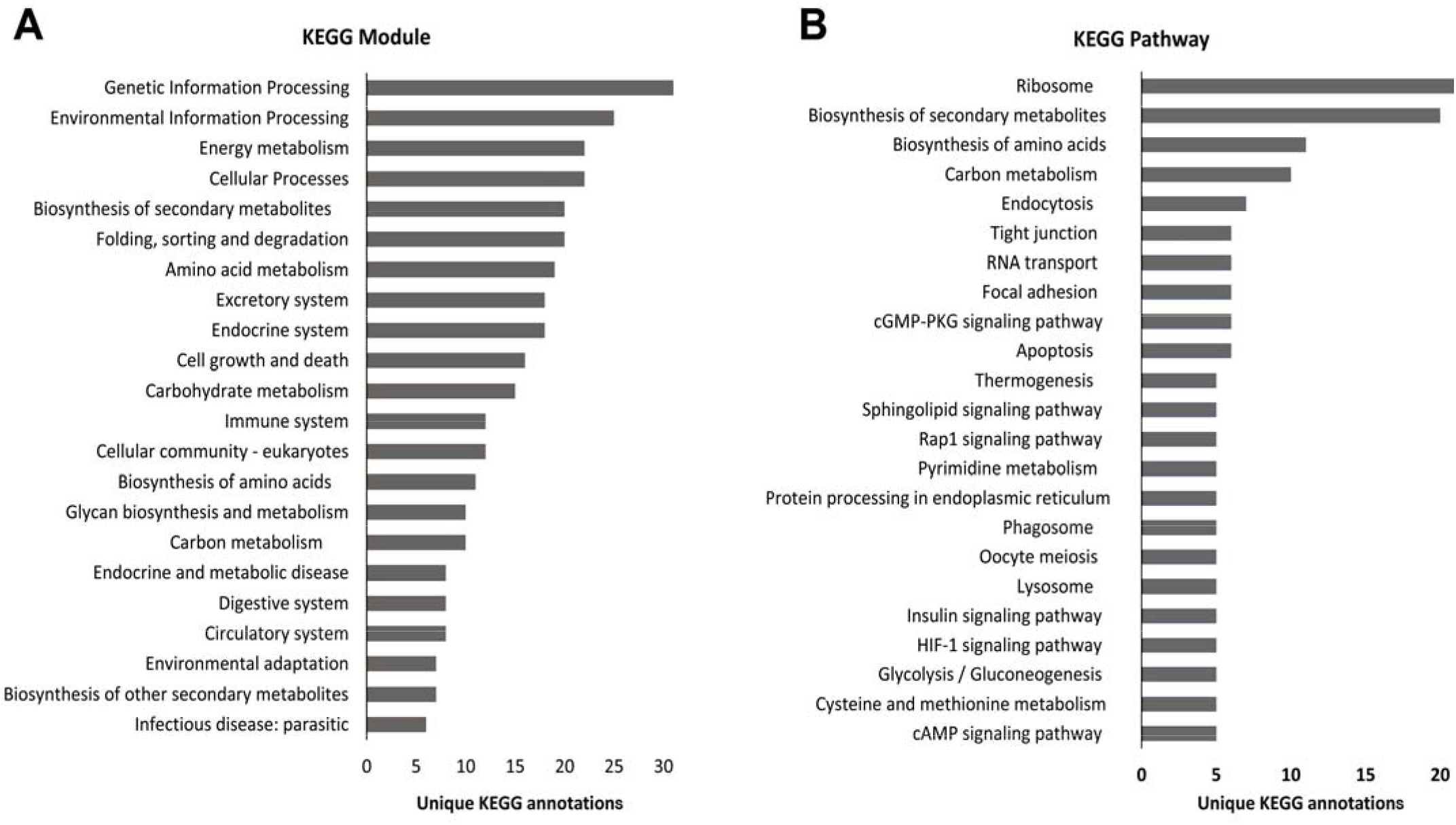
KEGG annotation of proteins, the levels of which significantly changed in worms that reached sexual maturity after incubation at 40 °C for 48 hours in comparison with infective plerocercoids. (A) Top KEGG modules. (B) Top KEGG pathways.

Only two proteins, arrestin A0A0X3NLP9 and uncharacterized protein A0A0X3PWQ6, were more abundant in plerocercoids than in adult worms (Tab. 1). At the transcriptome level, down-regulation of at least 5 arrestin isoforms in adult worms was previously detected by Hébert with colleagues (2017) (Table S4). Moreover, the maximum expression of the arrestin isoform A0A0X3NLP9 was recorded in pre-infective plerocercoids, which did not yet gain sufficient mass (<0.5 g), less in plerocercoids capable of infecting birds, and minimal in mature parasites, while the expression of the other 4 isoforms were maximal in infective plerocercoids. This indicates the great importance of arrestins, whose known function is signalling transduction, for the regulation of the second larval stage of *S. solidus*, although different isoforms regulate different periods of larval development. Sequence alignment of the uncharacterized protein A0A0X3PWQ6, suppressed in adult parasites, revealed orthologues in platyhelminths and a relationship with proteins of other organisms containing the Srp40 domain. This domain has been reported to be involved in ribosome processing, modification, and regulation of RNA polymerases; however, the significance of these proteins for flatworms remains to be elucidated.

Based on the analysis of the KEGG ontology (Kanehisa, 2019), the predominance of anabolic processes in the body of an adult worm can be stated, of which the pathways of protein synthesis and processing can be noted as the most affected (Fig. 3; Table S5). In mature worms, the level of 15 ribosome proteins, two proteins response for mRNA transport and splicing, one protein responsible for RNA degradation (RNA helicase DDX6), 5 translation initiators (components of translation initiation multifactor complex and eIF4G), one translation inhibitor (FMRP translational regulator factor) and two translation termination factors (eukaryotic peptide chain release factor, serine/threonine protein phosphatase 2A regulatory subunit) significantly increased (Figs S2, S3; Table S5). Signal peptidase complex subunit 3, two subunits of signal recognition particle (SRP) and four subunits of proteasome, which presumably determine post-translation fate of newly synthetized proteins, were also found to be abundant in the proteome of the mature parasite.

In adult worms, the increased expression of enzymes that regulate the metabolism of tyrosine, tryptophan, alanine, serine, glutathione, cysteine, glutamine, putrescine, spermidine, selenogomocysteine and others may indicate reinforced production of amino acids and regulators of protein synthesis. Glutamate dehydrogenase, a key nitrogen/carbon metabolism switching enzyme, was also elevated in mature worms. This enzyme provides the assimilation of ammonia in the form of glutamate or, conversely, releases α-ketoglutarate for the tricarboxylic acid cycle. Glutamate dehydrogenase has been found directly in trematodes vitellaria (Abidi et al., 2009) and its overexpression was observed in sexually mature parasites *S. solidus, Echinococcus granulosus, Opisthorchis felineus* and *Schistosoma mansoni* compared to their larvae and immature adults, indicating a crucial role in flatworm fertility (Protasio et al., 2012; Zheng et al., 2013; Hébert et al., 2017; Ershov et al., 2019). Interestingly, the *S. solidus* glutamate dehydrogenase isozyme A0A0X3P860 contains the sequence GFGNVG and Glu (278) in the structure, which is more typical for isoforms that provide the deamination reaction rather than the addition of ammonia (Oliveira et al., 2012; Prakash et al., 2018; Grzechowiak et al., 2020). Therefore, in adult flatworms, this enzyme appears to provide NADPH for redox reactions, probably for the malate dismutation pathway in mitochondria (Smyth and McManus, 2007; Ritler et al., 2019).

The utilization of energy reserves accumulated by the larva through glycogen breakdown initiated in adult *S. solidus* by the enhanced synthesis of glycogen phosphorylase A0A0V0J2F3 (Table 1) (Hopkins, 1952; Hébert et al., 2017). However, our study confirms that, in adult worms, the full set of glycolysis/gluconeogenesis enzymes is not activated, but only fructose-1,6-bisphosphatase, aldolase, glyceraldehyde-3-phosphate dehydrogenase and enolase (Table 1). Three out of five aldolase isoforms (A0A0X3PH87, A0A0X3Q357, A0A0X3NM70) identified in the mass spectra of the parasite were found only in adult worms. Moreover, enolases were not detected at all in the transcriptome and proteome of the plerocercoids, while in adults, 6 isoforms of the enzyme were detected by LC-MS/MS. Fructose-1,6-bisphosphatase A0A0X3NTV5, which catalyses the only irreversible reaction of gluconeogenesis, was also absent in the proteome of larvae and appeared in adult worms, while the concentration of fructokinases catalysing the competitive reverse reaction did not change. These results are in general consistent with Körting and Barrett study (1977), who reported the presence of low aldolase and fructose-1,6-bisphosphatase and enolase activity in *S. solidus* plerocercoids compared to other glycolytic enzymes.

Earlier, Hébert and colleagues (2017) recorded a decrease in the level of transcripts of other glycolytic enzymes in adult worms, but according to proteomic data, their content remains at the same level as in the larva. The activity of phosphoglycerate kinase, glucose-6-phosphate isomerase, phosphoglucomutase was reported to be quite high in *S. solidus* plerocercoids (Körting and Barrett, 1977), therefore glycolysis is possible in adult worms, but the carbon flux is most likely directed toward the production of glucose or glycolysis intermediates rather than the complete glucose breakdown (Fig. 4). We can also assume that overexpressed glycolytic enzymes play an alternative role in mature parasites, not only associated with the metabolism of their own carbohydrates. For example, they can be synthesized for export, since glycolysis enzymes have been found in the secret of the marita of the trematode *Opisthorchis felineus*; schistosomules of the trematode *Schistosoma japonicum*; *Taenia solium* metacestode and an adult *Hymenolepis diminuta* (Victor et al., 2012; Lvova et al., 2014; Bien et al., 2016; Cao et al., 2016). It is known that enolase is a marker of exosomes, vesicles that detach from the parasite’s cell membranes and deliver signalling molecules to their host (Buck et al., 2014; Samoil et al., 2018). In addition, various glycolytic enzymes are abundant in the shells and eggs of parasitic worms (Dewalick et al., 2011).

**Figure 4.**
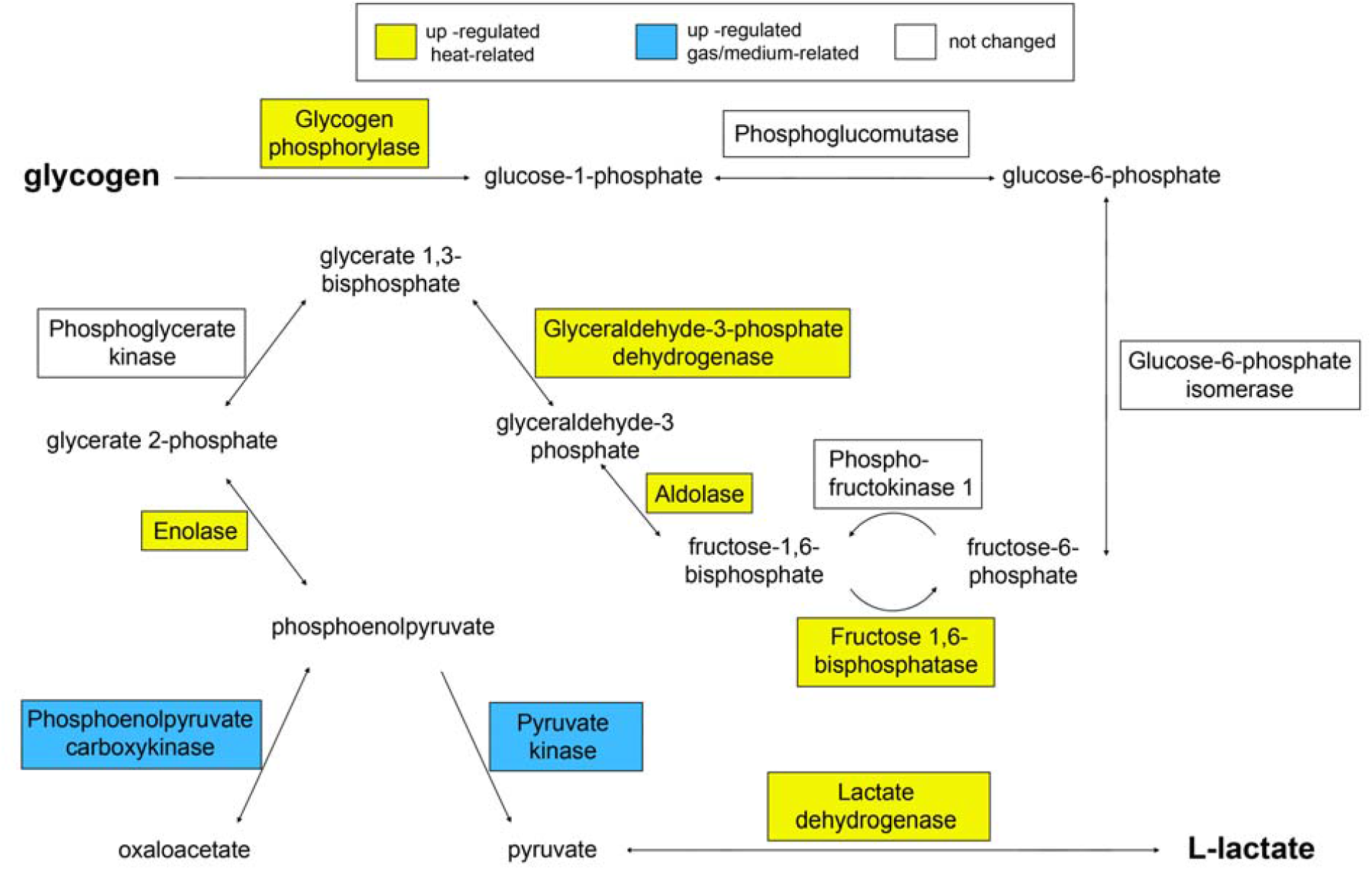
Scheme of glycolysis/gluconeogenesis pathway highlighting the proteins with increased concentration in adult worms compared with infective plerocercoids.

In cestodes, glucose oxidation leads either to the formation of lactate, or to the formation of oxaloacetate, and then malate, which is converted into succinate and acetate in mitochondria (CO_2_ fixation pathway; malate dismutation pathway) (Smyth and McManus, 2007; Ritler et al, 2019). This cleavage occurs at the level of phosphoenolpyruvate, for which pyruvate kinase (PK) and phosphoenolpyruvate carboxykinase (PEPCK) compete (Das et al., 2015) (Fig. 4). The level of pyruvate kinase A0A0X3Q057 and one of the PEPCK isoforms A0A183TAB4 was increased in worms kept under anaerobic conditions at both 40 °C and 22 °C, which suggests that the expression of these enzymes can be controlled by the gas composition of the medium. In turn, pyruvate formed by pyruvate kinase is converted by lactate dehydrogenase to lactate, which is the end product of *S. solidus* metabolism in a warm-blooded host (Smyth and McManus, 2007; Beis and Barrett, 1979). We found that two of the three isoforms of lactate dehydrogenases (A0A183TTF7, A0A3P7E2D4) identified in the mass spectra of the parasite were present only in adult worm proteomes. This enzyme promotes energy production when there is a lack of oxygen in the intestines of birds; however, in our experiment, adult-specific isoforms were not detected in extracts of worms incubated under semi-anaerobic conditions at 22 °C. Our study confirms the conclusion of Hébert and colleagues (2017) that the synthesis of enolase and some isoforms of aldolase, fructose-1,6-bisphosphatase and lactate dehydrogenase is a phenotypic marker of the sexually mature stage of *S. solidus* (ecological status of the annotation “switched ON in the bird”), since these enzymes are under the control of genes, which are induced by an increase in temperature. In turn, the fate of the glycolysis product, phosphoenolpyruvate, is controlled by genes regulated by other factors, most likely oxygen or carbon dioxide.

It was expected that the synthesis of proteins that regulate reproductive function and immune response would be altered in gravid *S. solidus* compared to invective plerocercoids obtained from the fish body cavity. Among the heat-induced proteins that are potentially involved in the reproduction process in *S. solidus*, mitogen-activated protein kinase A0A0X3PWN3 can be noted, which is one of the key enzymes of oocyte meiosis and fertilization (Fan and Sun, 2004). Many of *S. solidus* protein kinases and protein phosphatases activated by heat can also participate in gametogenesis and impregnation in gravid worms by coordinating the activity of A0A0X3PWN3 (Table S5). Further studies of the localization of these proteins in adult worms are necessary to clarify their significance for the reproduction of cestodes.

At the proteomic level, we did not observe a significant parasitic immune evasion of adult *S. solidus* compared to infective plerocercoids. Only 12 proteins, elevated by incubation at high temperature, were KEGG annotated as potentially involved in the regulation of the immune system. Of these, 7 are nonspecific regulators of signal transduction (protein kinases A0A0X3PXQ5, A0A0X3PWN3, A0A0X3PTZ8, phosphatase A0A0X3NTL5, Ras proteins A0A183SMP0, A0A0X3Q3W4 and 14-3-3 protein A0A0V0J2Z1), 2 involved in protein folding (calnexin A0A0X3NKP9 and peptidyl-prolyl cis-trans isomerase A0A3P7CFD4), and 2 associated with cell motility and adhesion (actin A0A0X3PZ06 and vinculin A0A0X3P131). Only the immunomodulatory cathepsin B-like peptidase A0A183SQT2 had a signal peptide for secretion and, more likely, could be involved in the regulation of final host immunity. This protein has been found in excretory/secretory products of *Echinococcus multilocularis* and gut extracts of parasitic nematode, and has been shown to degrade host connective tissue, suggesting an important role in helminth digestion and invasion (Sako et al., 2011; Long et al., 2015; Caffrey et al., 2018). Since adult *S. solidus* have impaired nutrition (Hopkins, 1950) and do not penetrate the avian intestinal wall, it can be assumed that the main reason for the expression of this protein in adult worms is the ability to destroy host immunoglobulins and components of the compliment (Sako et al., 2011). Summarizing, it seems that anti-immune response of the parasite in the final host is not controlled by temperature; however, the possibility of inducing parasite defence by direct interaction with host immune cells remains questionable.

## Conclusions

In this study, we analysed of the spectra of *S. solidus* proteins to explore the molecular adaptations of parasites during trophic transmission from cold-blooded fish to warm-blooded final hosts. To identify the initial events that occur in a warm-blooded host, we studied changes in the spectrum of *S. solidus* proteins in the first hour after heating and found structural rearrangements of the parasite tegument at the molecular level. The long-term changes in warm-blooded host are primarily associated with the consumption of energy stored by the larvae in the form of glycogen for protein biosynthesis. The observed activation of amino acids metabolism and protein production pathways in gravid worms presumably enable gametogenesis and egg production. Comparison of the obtained data with the results of studies analysing the transcriptome of this parasite confirmed the previously described changes in *S. solidus* gene expression in the final host, as well as revealed new markers of certain stages of worm development. We believe that further studies of the localization of stage-specific helminth proteins will clarify their role in maintaining the parasitic lifestyle and in the relationship with hosts.

## Acknowledgements

We express our gratitude to the members of the Department of Ichthyology and Hydrobiology of Saint Petersburg State University, PhD Dmitry L. Lajus, PhD Mihail V. Ivanov and Tat’yana S. Ivanova for assistance in obtaining biological material. The LC-MS/MS analysis was performed using equipment of ’Human proteome” Core Facility of the Institute of Biomedical Chemistry (IBMC, Moscow, Russia). The equipment of the Core Facility of the Karelian Research Centre of the Russian Academy of Sciences was used during animal experiment and protein purification.

## Data availability

The mass spectrometry proteomics data have been deposited to the ProteomeXchange Consortium via the PRIDE [1] partner repository with the dataset identifier PXD024166”. Project Name: Proteome of *Schistocephalus solidus* (Cestoda) after short- and long-term heating.

